# Fungal polysaccharopeptides reduce obesity by richness of specific microbiota and modulation of lipid metabolism

**DOI:** 10.1101/375584

**Authors:** Xiaojun Li, Peng Chen, Peng Zhang, Yifan Chang, Mingxu Cui, Jinyou Duan

**Author notes:** Correspondence: Jinyou Duan, PhD.

## Abstract

The prevalence of obesity and related disorders has vastly increased throughout the world and prevention of such circumstances thus represents a major challenge. Here, we show that protein-bound β-glucan (PBG), one representative of *Coriolus versicolor* polysaccharopeptides which are broadly used as immune boosters and clinically implicated in treatment of cancers and chronic hepatitis, could be a potent anti-obesity agent. PBG could reduce obesity and metabolic inflammation in mice fed with a high-fat diet (HFD). Gut microbiota analysis revealed that PBG markedly increased the abundance of *Akkermansia muciniphila* although it didn’t rescue HFD-induced change in the *Firmicutes* to *Bacteroidetes* ratio. It appeared that PBG altered host physiology and created an intestinal microenvironment favorable for *A. muciniphila* colonization. Fecal transplants from PBG-treated animals in part reduced obesity in recipient HFD-fed mice. Further, PBG was shown to promote lipid metabolism in microbiota-depleted mice. Thus, our data highlights that PBG might exert its anti-obesity effects through a mirobiota-dependent (richness of specific microbiota) and -independent (modulation of lipid metabolism) manner. The fact that *Coriolus versicolor* polysaccharopeptides are approved oral immune boosters in cancers and chronic hepatitis with well-established safety profiles may accelerate the development of PBG as a novel drug for obesity treatment.

## Introduction

Obesity, an abnormal or excessive fat accumulation in adipose tissues, has become a major public health issue worldwide. Obesity is well-known as a contributing factor in diabetes, cardiovascular disease, hypertension, stroke, and cancer, etc (1). Given the serious health consequences of obesity as well as its economic burden, greater attention must be directed to the prevention, identification, and treatment of overweight and obese conditions (2).

Despite the fact that dieting and physical exercise is the main treatment modalities, persons with obesity failing lifestyle therapies need further aggressive intervention including pharmacotherapy, medical devices and bariatric surgery (3–5). Noninvasive anti-obesity drugs have resurfaced as adjunctive therapeutic approaches to bridge a gap between intensive lifestyle intervention and surgery (6).

Drug repurposing, also known as drug repositioning, is a strategy that seeks to reuse existing, licensed medications for new medical indications (7). It offers a variety of advantages when compared with *de novo* drug development. These include the availability of extensive pharmacokinetic, pharmacodynamic and safety data, and an understanding of relevant molecular targets and mechanisms of action, due to previous considerable clinical experience (8). Thus, drug repurposing can greatly slash development costs and quickly translate those existing medications into treatment of epidemic or devastating diseases (9).

Many cultures worldwide have long recognized that hot water decoctions from certain medicinal mushrooms such as *Ganoderma lucidum* and *Coriolus versicolor* have health-promoting benefits (10). Thereinto, the polysaccharide fractions are believed as the major immunotherapeutic components, which are either β-glucans, β-glucans with heterosaccharide chains or protein-bound β-glucans (PBGs), in other words, polysaccharopeptides (11–13). Numerous investigations *in vitro, in vivo* and clinical trials have reported that crude or purified PBGs from *C. versicolor* were adopted as an adjunct therapy for cancer (14) or chronic hepatitis (15). In this study, one purified PBG was repositioning as a potent anti-obesity agent and the in-depth understanding of this beneficial effect was examined.

## Results

### PBG prevents HFD-induced obesity

Oral administration of PBG daily at either a low dose (200 mg/kg, Group L) or a high dose (400 mg/kg, Group H) leads to a significant decrease of body weight in HFD-fed mice from 5 to 9 weeks (Fig.1a). The decreased weight gain (Fig.1b) and liver weight (Fig.1c) in PBG-treated mice at 9 weeks is not due to reduced food consumption (Fig.1d). PBG substantially prevents HFD-induced fat accumulation (Fig.1e), while it has no effect on lean mass (Fig.1f). Magnetic resonance imaging (MRI) reveals that PBG treatment apparently attenuates fat area in HFD-fed mice (Fig.1gh). Consistent with macroscopic evaluation of adiposity above, PBG feeding results in a dramatic reduction of triglyceride contents (Fig.1i), total bile acid (Fig.1j) and leptin (Fig.1k) in serum. There is an increase of plasma adiponectin (Fig.1l) and a decrease of hepatic lipid droplets (Fig.1mn) in HFD-fed mice after PBG treatment. It is noted that PBG daily at a dose of 200 mg/kg elicits an anti-obesity effect, comparable to simvastatin (2 mg/kg), a lipid-lowering medication on the World Health Organization’s List of Essential Medicines while it is believed to be contraindicated with pregnancy, breast feeding, and liver disease (16).

**Fig.1.**
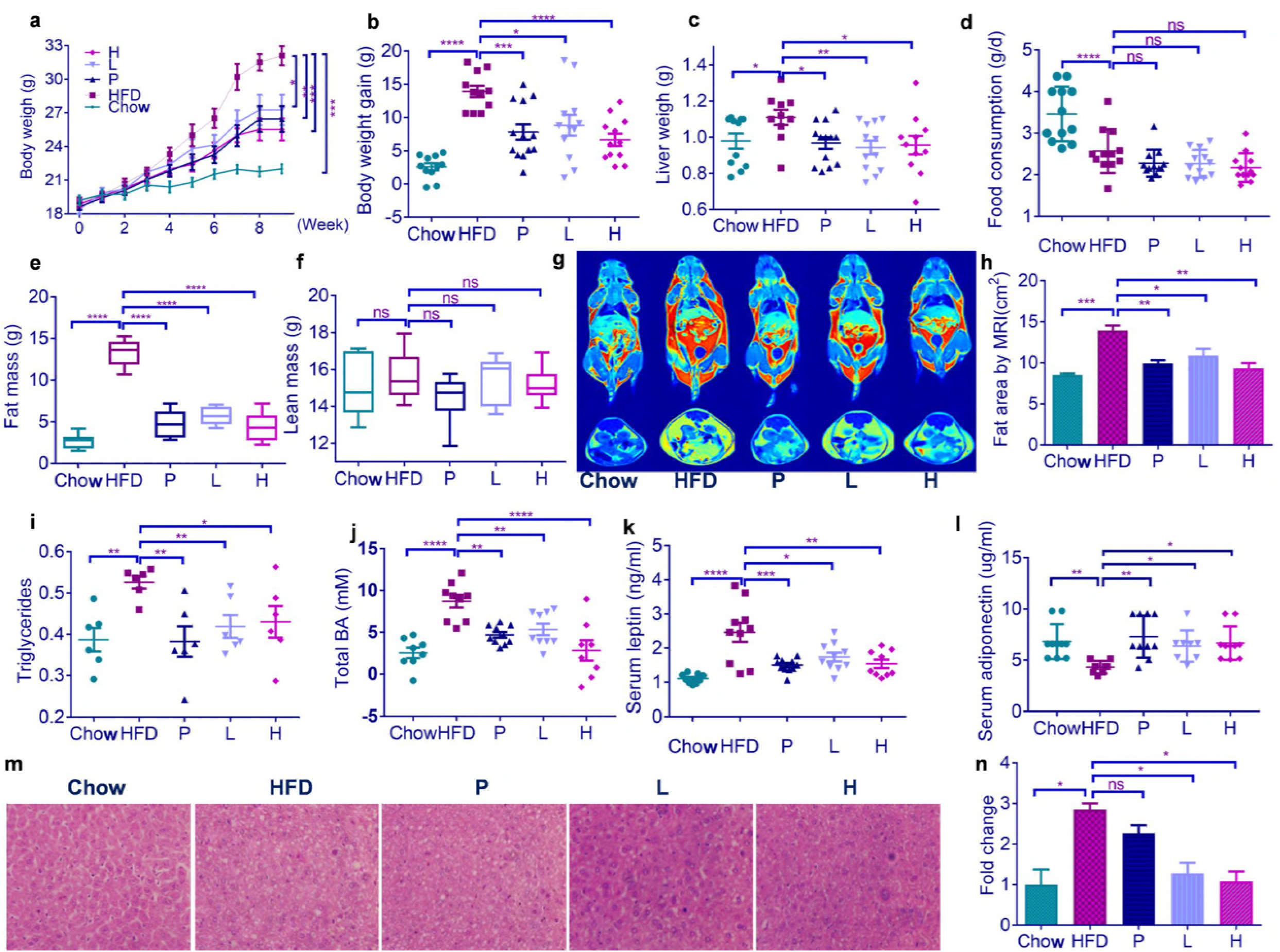
PBG treatment decreases body weight and fat accumulation in HFD-induced obese mice. HFD-fed mice were treated daily with either water (group HFD), simvastatin at 2 mg/kg (Group P) or PBG at 200 (Group L) or 400 mg/kg (Group H) by intragastric gavage for 9 weeks (n=11-13 for each group). As a control, Chow-fed mice (Group Chow) were lavaged with water daily. (a) body weight curves; (b) body weight gain; (c) liver weight; (d) food consumption; (e) fat mass; (f) lean mass; (g) a representative of magnetic resonance imaging (MRI); (h) fat area by MRI; (i) serum triglyceride; (j) total bile acid in serum; (k) serum leptin; (l) serum adiponectin; (m) a representative of liver sections in which lipid content was assessed using H&E staining; (h) normalized fold change of liver lipid content. Data are shown as mean±s.e.m. *p* value in (a) and (l) were analysed using unpaired two-tailed Student’s t-test. *p* value in (b)-(k) and (n) were analysed using Dunnett’s multiple comparisons test. \**p* < 0.05, \*\**p* < 0.01, \*\*\**p* < 0.001 and \*\*\*\**p* < 0.0001.

### PBG reduces obesity-related metabolic inflammation

Obesity is always associated with a low-grade chronic inflammation (17). HFD-fed mice produce higher levels of pro-inflammatory cytokines including tumors necrosis factor-alpha (TNF-α), interleukin-1-beta (IL-1β), interleukin-17A (IL-17A) and interleukin-6 (IL-6) in serum than chow-fed mice do (Fig.2a-d), indicating the systemic inflammation is induced by HFD feeding. Administration of PBG or simvastatin suppresses the overproduction of those pro-inflammatory cytokines above in HFD-fed mice.

**Fig.2.**
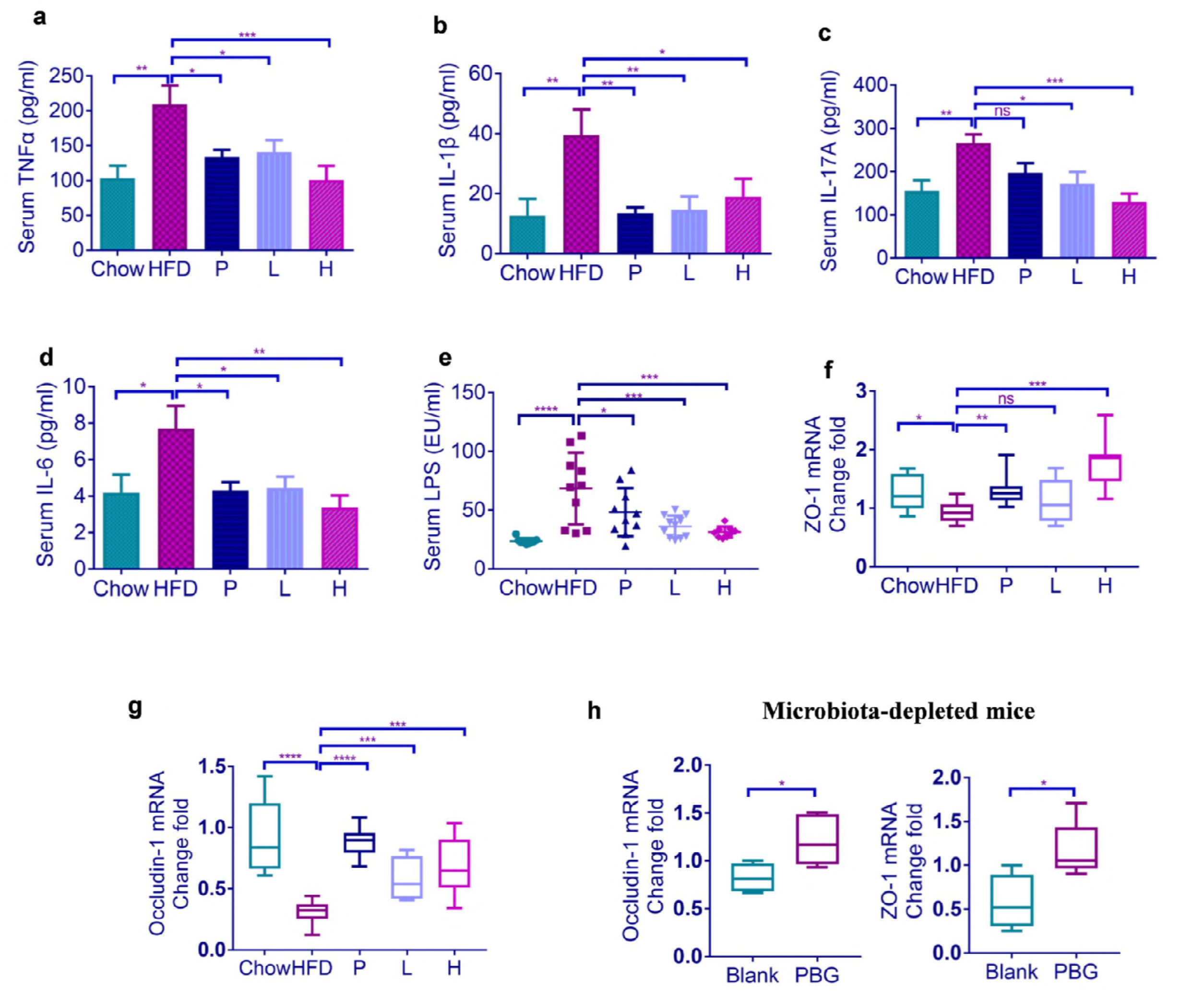
PBG treatment reduces serum pro-inflammation cytokines and lipopolysaccharide through improvement of intestinal barrier integrity in obese mice. Mice were treated as Figure 1. Effects of PBG treatment on serum TNF-α, IL-1β, IL-17A, IL-6 and endotoxin (a-e) were examined by ELISA kits, and on relative mRNA expression of ZO-1(f) and occludin (g) in colon were assessed using qRT-PCR; (h) relative mRNA expression of ZO-1 and occludin from colons of mirobiota-depleted mice treated with or without PBG. *p* value in (a-e) were analysed using Dunnett’s multiple comparisons test. *p* value in (f-h) were analyzed using unpaired two-tailed Student’s t-test. \**p* < 0.05, \*\**p* < 0.01, \*\*\**p* < 0.001,\*\*\*\**p* < 0.0001.

The obesity-related inflammation is characterized with an elevated level of gut microbiota-derived lipopolysaccharide (LPS) in circulation, an event defined as metabolic endotoxemia (18). PBG or simvastatin significantly reduces serum LPS levels in HFD-fed mice (Fig.2e), and increases expression of the tight junction components, zonula occludens-1 (ZO-1) (Fig.2f) and occludin-1 (Fig.2g). It seems that PBG can upregulate the transcription of ZO-1 and occluding-1 in microbiota-depleted mice, suggesting PBG directly protects HFD-induced disruption of tight junction regardless of gut microbiota (Fig.2h). The findings above are consistent with the concept that the increased expression of intestinal tight junction proteins improves intestinal barrier integrity and thus decreases leakage of microbial LPS from gut into circulation (17).

### PBG beneficially alters obese-type gut microbiota

It is generally accepted that changes in the composition of gut microbiota are associated with the development of obesity (19). To assess whether gut microbiota is involved in the anti-obesity effects of PBG, we performed a pyrosequencing-based assay of bacterial 16S rRNA (V3-V4 region) in fecal samples from different groups of mice (11-13 mice/group). After removing unqualified sequences (see Methods), a total of 4,799,464 raw reads and an average of 67,738±3,949 reads per sample were obtained. After selecting the effective reads, a total of 3,244,442 effective reads was generated and each fecal sample produced an average of 54,074±4,309 effective reads. Samples with a low number of effective reads (<1622) were not observed. High-quality reads were clustered into Operational Taxonomic Units (OTUs).

Principal coordinates analysis (PCoA) of unweighted UniFrac distances of fecal microbiota was plotted, based on OTU abundances (Fig. 3a). The PCoA scores clearly display a distinct clustering of microbiota composition for each treatment group. Simvastatin treatment induces a mild shift of gut microbiota in HFD-fed mice, while intake of PBG leads to a more pronounced change in microbiota composition. The UPGMA (unweighted pair-group method with arithmetic means) tree also indicates there is a remarkable separation between the microbiota from most mice for each treatment (Fig. 3b).

**Fig.3.**
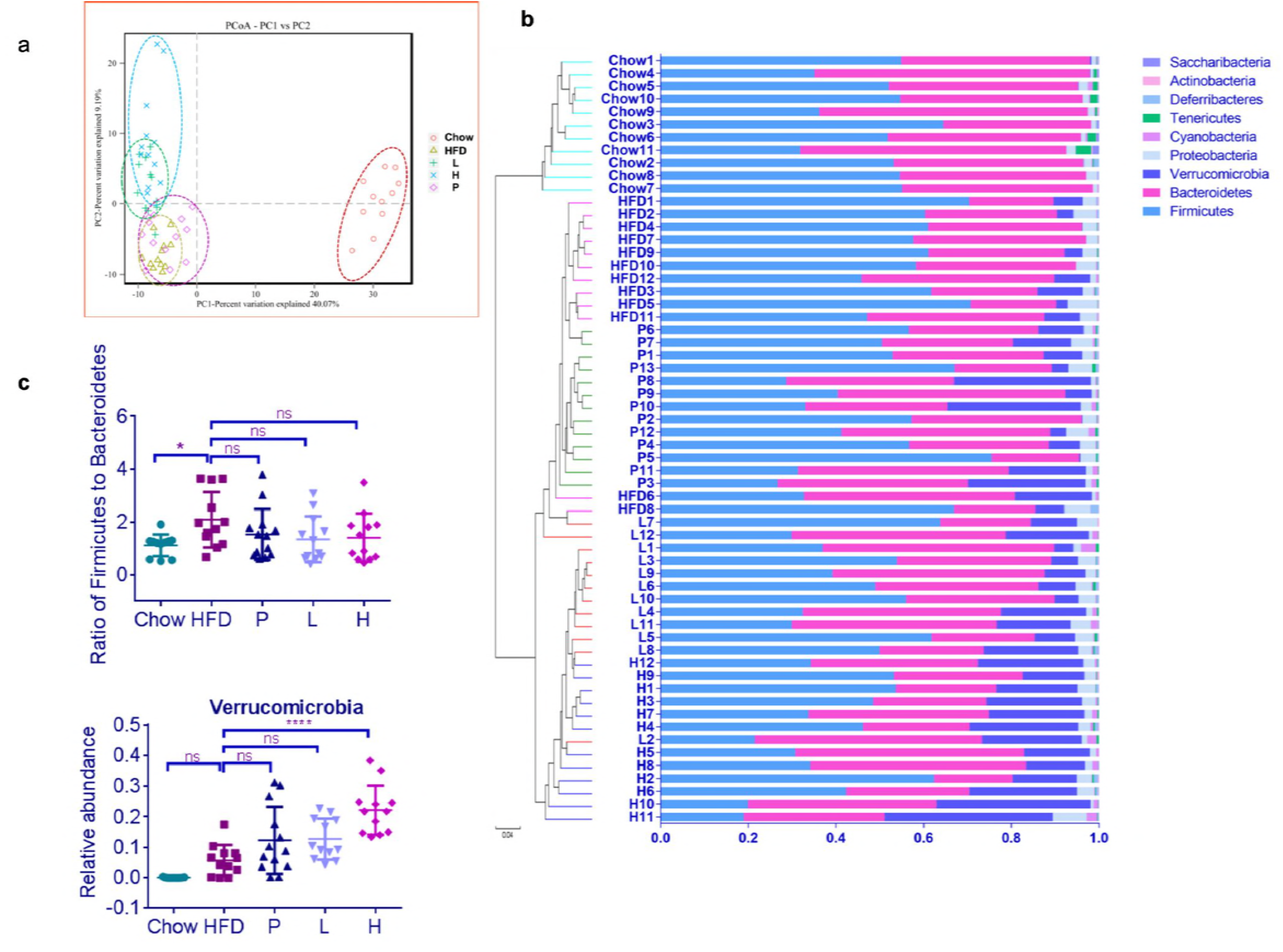
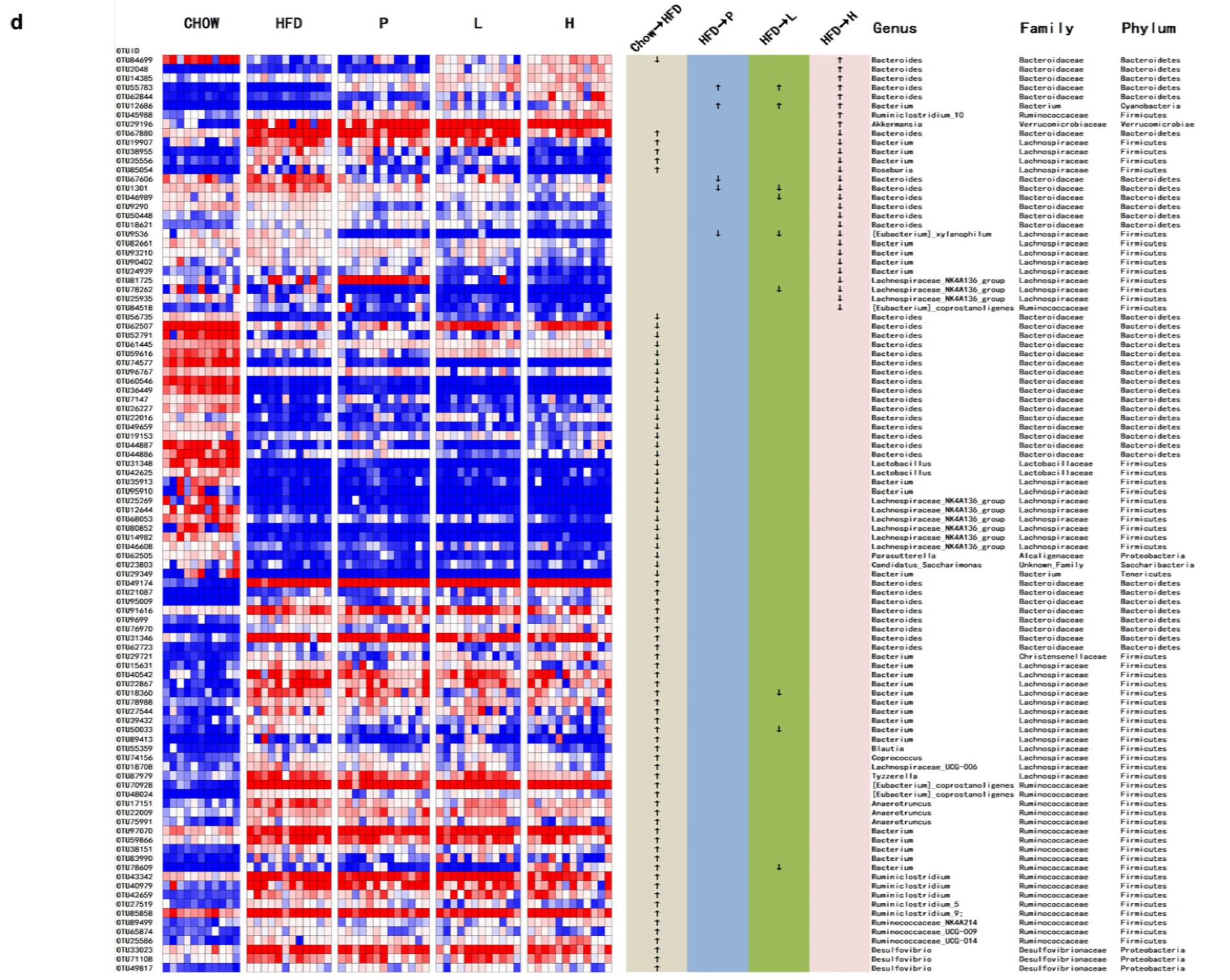
PBG treatment affects mcirobiota in HFD-induced obese mice. Microbiota composition were analyzed using a pyrosequencing-based assay on feces of chow- and HFD-fed mice (n=11-13 for each group) as described in Figure 1. (a) Plots were generated using the unweighted version of the UniFrac-based PcoA; (b) Bacterial taxonomic profiling in the phylum level of microbiota from individual groups; (c) Comparison of *Firmicutes* to *Bacteroidetes* ratio and relative abundance of the phylm *Verrucomicrobia. p* values were analyzed using Dunnett’s multiple comparisons test (\**p* < 0.05, \*\**p* < 0.01, \*\*\**p* < 0.001, \*\*\*\**p* < 0.0001). (d) Heatmap and bacterial taxa information (genus, family and phylum) of the Top 100 OTUs. Difference was analyzed using Paired Wilcoxon rank-sum test (\**p* < 0.05).

Analysis of bacterial relative abundance confirmed prior reports that gut microbial communities of mice are dominated by bacteria from the *Firmicutes* and *Bacteroidetes* phyla (Fig. 3b) and that HFD feeding increases the proportion of *Firmicutes* versus *Bacteroidetes* (20). It appear that either PBG or simvastatin treatment doesn’t rescue HFD-induced change in the *Firmicutes* to *Bacteroidetes* ratio (Fig. 3c). The most striking difference in gut microbiota between HFD-fed mice and PBG-treated mice lies in the abundance of the phylm *Verrucomicrobia* (Fig. 3c).

We used Wilcoxon signed-rank test to identify the specific bacterial genera that were altered by HFD feeding and PBG treatment (Fig. 3d). Compared with chow-fed mice, HFD feeding significantly altered 420 OTUs, producing 249 increased and 171 decreased OTUs. In HFD-fed mice, simvastatin treatment altered 36 OTUs (13 increased and 23 decreased), while PGB daily at a dose of 400 mg/kg altered 140 OTUs (29 increased and 111 decreased). Detailed analysis of the Top 100 OTUs indicated that the following 4 genera, *Akkermansia, Bacteroides, Bacterium*, and *Ruminiclostridium_10* were profoundly increased in PBG-treated HFD-fed mice. The most striking one was *Akkermansia*, which was identified as the sole genus in the phylm *Verrucomicrobia*. PBG increased the percentage of *Akkermansia* from 5.67 ± 1.5% to 22.1 ± 2.3% (*p* < 0.0001) at a daily dose of 400 mg/kg.

### PBG promotes *Akkermansia muciniphila* colonization through altered host physiology

It remains to be determined whether the interaction between PBG and the gut microbiota, *Akkermansia* is direct or indirect (i.e., mediated through promoting bacterial growth or altering host physiology). The *in vitro* analysis indicated that PBG couldn’t directly promote the growth of *A. muciniphila* in the pure culture (Fig.4a). To see whether PBG affected the growth of *A. muciniphila* in a complex microbiota, we cultured fecal samples supplemented with *A. muciniphila* in two separate gut-simulators, and exposed them to a constant flow of PBG (40 mg/mL) for 42 h. No substantial difference in the abundance of *A. muciniphila* was observed following PBG exposure (Fig.4b).

**Fig.4.**
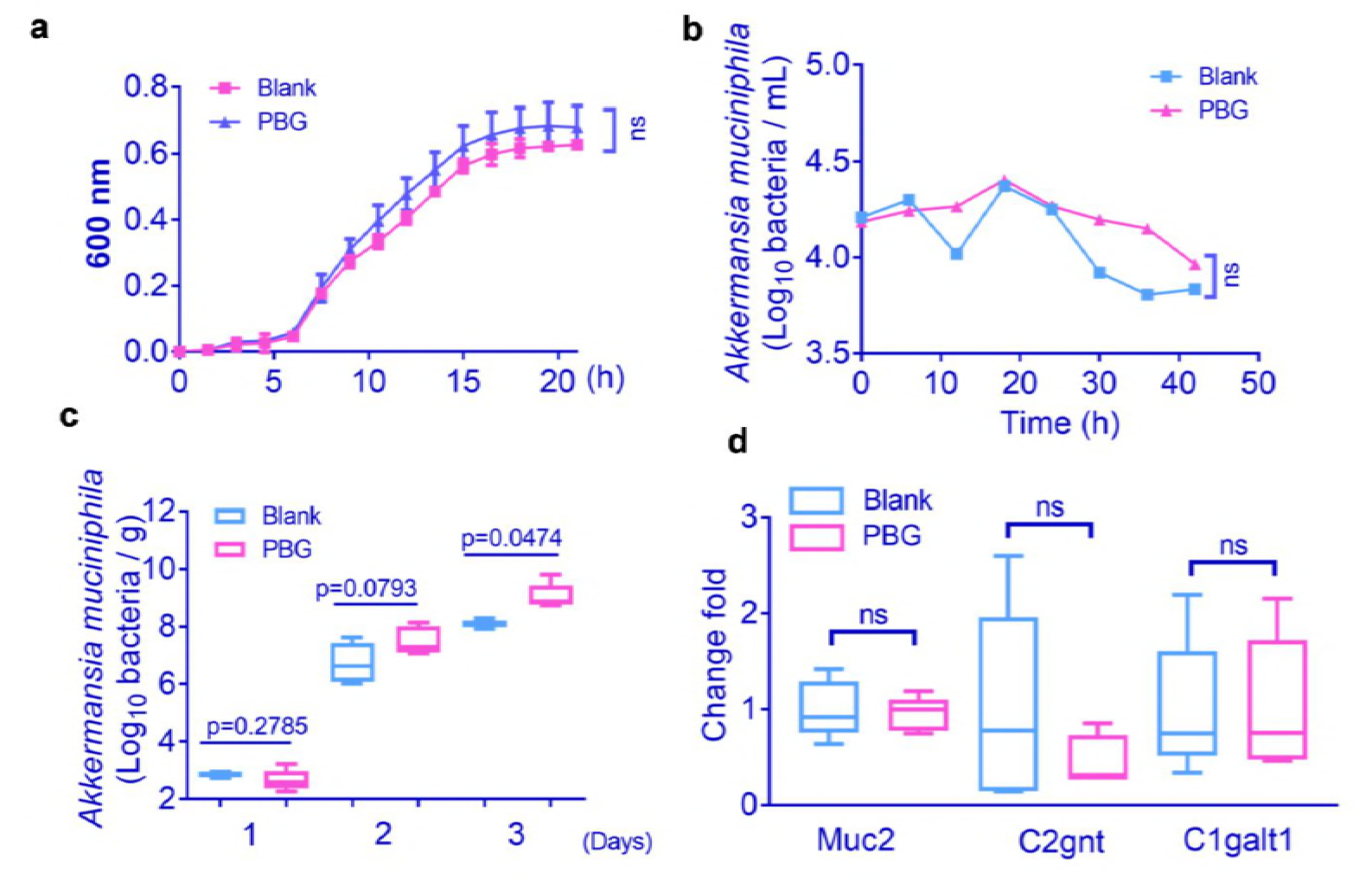
Effects of PBG on *A. muciniphila* growth and colonization as well as gene expression of intestinal mucin 2. (a) Growth of *A. muciniphila* as a single culture in the presence or absence of 10-mM PBG (with six technical replicates); (b) Growth of *A. muciniphila* in complex microbiota using the *in vitro* gut simulator with or without PBG exposure; (c) Effects of PBG treatment on *A. muciniphila* colonization in microbiota-depleted mice. HFD-fed mice were treated daily with a cocktail of antibiotics for one month, followed by supplementation with or without PBG (400 mg/kg) for another month in the presence of antibiotics. Mice were lavaged with *A. muciniphila* (1.5 × 10^8^ cfu) continuously for 3 days. *A. muciniphila* in the fecal samples was determined 24 h after each lavage; (d) Effects of PBG treatment on gene expression of protein and glycosyltransferases of mucin 2 in the colon tissues from microbiota-depleted mice. *p* value was analyzed using unpaired two-tailed Student’s t-test (\**p* < 0.05).

To address whether PBG could alter host physiology and thus facilitate *A. muciniphila* colonization, we treated mice with an antibiotic cocktail for one month to deplete gut microbiota. The microbiota-depleted mice were further treated daily with PBG or PBS, along with the antibiotic cocktail to prevent recurrence of microbiota for another month. The mice were lavaged with *A. muciniphila* (2 × 10^8^ cfu / 0.2ml) continuously for 3 days and its abundance in the fecal samples was determined 24 h after each lavage. The amount of *A. muciniphila* was statically different at day 3 in feces between PBG-treated mice and control mice (Fig.4c). This observation suggested that PBG created an intestinal microenvironment favorable for *A. muciniphila* colonization.

*A. muciniphila* degrades intestinal mucins, the highly glycosylated proteins of the epithelial mucus layer, as its preferred source of carbon and nitrogen (21). Alterations of the mucus could be caused either by modification of gene expression that encode for the mucins (MUC genes) and/or of genes encoding for glycosyltransferases (GT genes) (22–24). We did not observe significant differences in gene expression of mucin 2 protein (encode by MUC2 gene) and its related glycosyltransferases (C1galt1 and C2gnt genes) in colon tissues from PBG-treated mice and control mice (Fig.4d), suggesting that *A. muciniphila* is not simply responding to increased host mucin production.

### Fecal microbiota transplantation from PBG-treated mice reduces obesity

It is well-known that alteration of gut microbiota by the transfer of foreign fecal materials could modulate obesity (25). We daily transferred fecal microbiota from the donor mice including chow-fed one, PBG treated chow-fed one, PBG treated HFD-fed one and HFD-fed one to the recipient HFD-fed mice. After 8 weeks, obesity-related syndromes were examined. The fecal transfer from the donor mice fed with chow (Chow→HFD), chow and PBG [PBG(Chow)→HFD], or HFD and PBG [PBG (HFD)→HFD] reduced body weight, plasma leptin and total bile acid and increased plasma adiponectin, compared with that from HFD-fed mice (HFD→HFD) (Fig. 5a–c). Correspondingly, the obesity-related metabolic inflammation, as indicated by elevated serum levels of proinflammatory cytokines such as IL-6, IL-1β and TNF-α in the recipient mice was also diminished after the first three fecal microbiota transplantation above (Fig. 5d–f). These results suggest that the anti-obesity effects of PBG in HFD-fed mice might be due to modulation of the gut microbiota.

**Fig.5.**
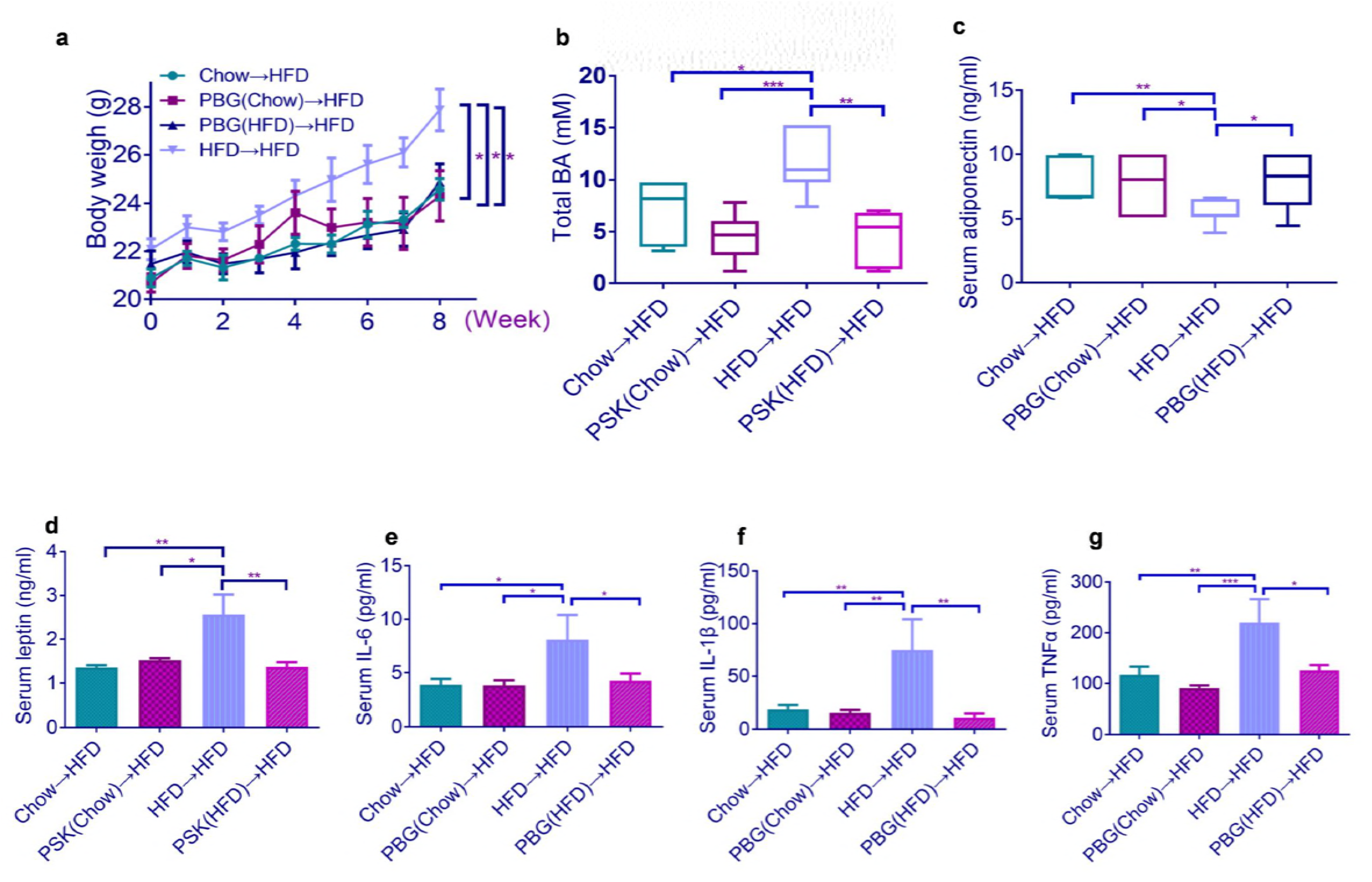
Obesity and related metabolic inflammation were reversed by fecal transplantation from PBG-treated mice to HFD-fed mice. HFD-fed mice were colonized with feces from different mouse groups for 8 weeks (n=6-8 for each group), followed by measurement of body weight (a), total bile acid (b), serum adiponectin (c), leptin (d), IL-6 (e), IL-1β (f), and TNF-α (g). *p* value in (a) and (c) were analyzed using unpaired two-tailed Student’s t-test. *p* value in (b) and (d)-(g) were analyzed using Dunnett’s multiple comparisons test. \**p* < 0.05, \*\**p* < 0.01, \*\*\**p* < 0.001,\*\*\*\**p* < 0.0001.

### PBG directly upregulates a set of gene expression involved in host metabolism

To find out whether PBG has a direct impact on host metabolism, we performed transcriptomic analysis of colon tissues from microbiota-depleted mice treated with or without PBG. After quality control, a total of 130,093,406 clean reads and 38.78 Gb clean bases were obtained. The percentage of Q30 bases in each sample was not less than 90.94%. A total of 22,419 UniGenes were identified. To facilitate the functional analysis of these RNAs, the differential expressed mRNA genes were picked from the whole gene matrix with Cuffdiff (26) according to the two standards: |log2(Fold Change)| ≧ 1 and adjusted *p*-value < 0.05. Compared with the control group, the PBG group had 155 differential expressed genes (DEGs) (Fig. 6a). Among 155 DEGs, 120 were successfully annotated by Gene Ontology (GO) assignments and classified into three functional categories, which were molecular function, biological process, and cellular component. There were less DEGs classified into cellular component than the other two categories (Fig. 6b).

**Fig.6.**
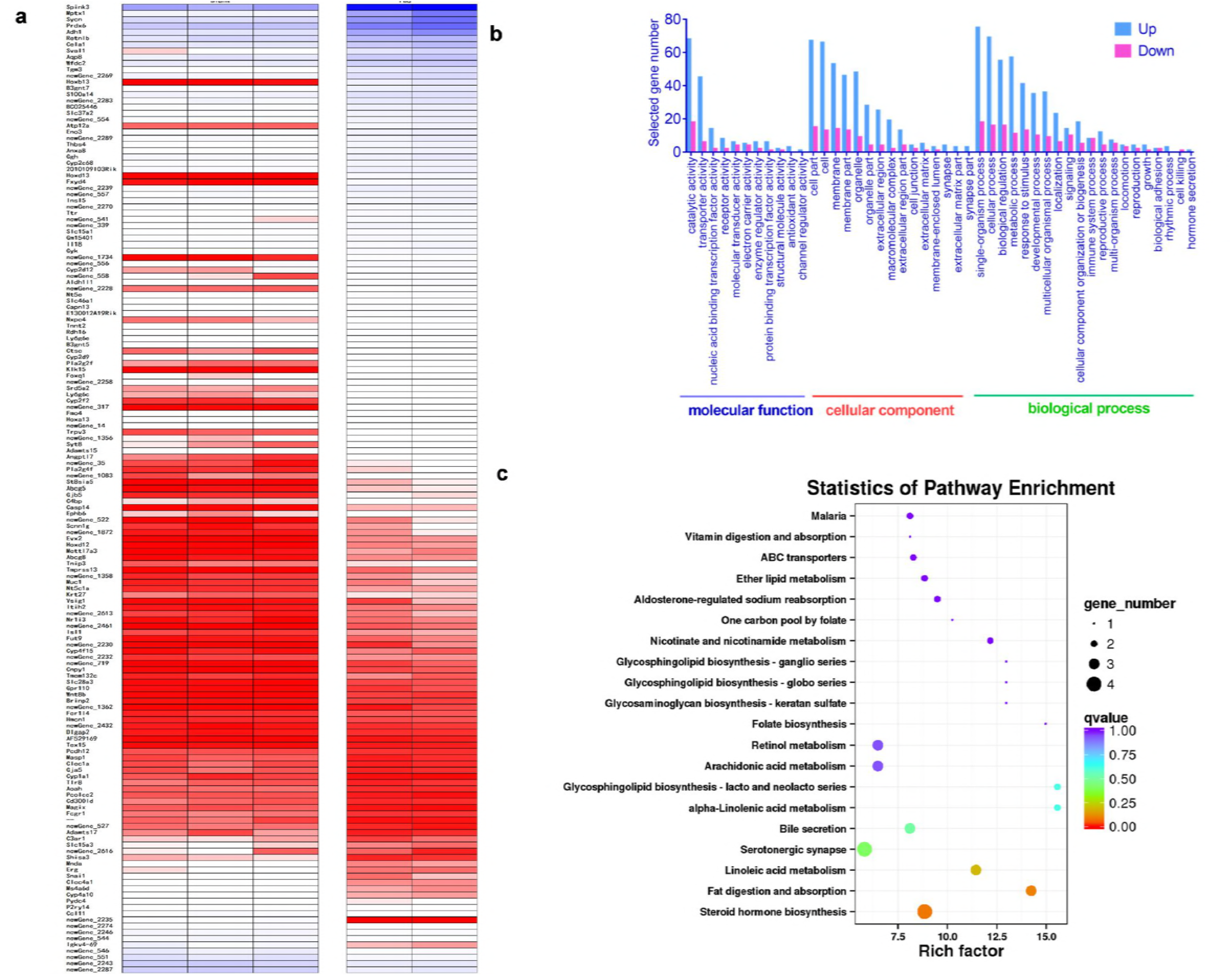
Transcriptome analysis of differentially expressed genes (DEGs) in colon tissues of microbiota-depleted mice treated with or without PBG. HFD-fed mice were treated daily with a cocktail of antibiotics for one month, followed by supplementation with or without PBG (400 mg/kg) for another month in the presence of antibiotics. (a) heatmap of total 155 DEGs; (b) Gene Ontology (GO) assignments of DEGs; (c) KEGG enrichment analysis of PBG-induced upregulated DEGs. The top 20 pathways were shown. The x-axis indicates the enrichment factor, and the y-axis shows the KEGG pathway.

Within 155 DEGs, KEGG (Kyoto Encyclopedia of Genes and Genomes) database analysis revealed 24 DEGs were relevant to metabolic pathways. The pathways related to lipid metabolism were mostly enriched, including fat digestion and absorption, glycosphinglipid biosynthesis, linoleic aicd metabolism and bile secretion (Fig. 6c). These included a set of DEGs such as B3gnt5, Abcg8, Gyk, St8sia5, Pla2g2f, Aqp8, Abcg5, Pla2g4f, Fut9, Adh1, and Cyp2c68 which were highly upregulated in colon tissues from PBG-treated mice (Table S1). These findings indicated that PBG could promote host lipid metabolism, regardless of gut microbiota.

## Discussion

The polysaccharopeptides produced by *C. versicolor* are effective immunopotentiators, supplementing the chemotherapy and radiotherapy of caners and infectious diseases. Several kinds of polysaccharopeptides have been shown to be produced by cultured mycelia or fruiting bodies. Although some of these polymers are structurally distinct, they are not distinguishable in terms of physiological activities (27).

PBG, one representative of polysaccharopeptides from the fruiting bodies of *C. versicolor*, indeed upregulates expression of genes related to host immune response in colon tissues (Table S1). The most pronounced one is CTSE gene, which encodes cathepsin E, an aspartic endopeptidase involved in antigen processing via the MHC class II pathway (28). Besides, several sets of genes related to elements of Toll-like or NOD-like signaling pathway and the complement system are upregulated (Table S1). These data suggest that PBG can activate innate and adaptive immunity in the intestine. Paradoxically, PBG suppresses systemic metabolic inflammation in HFD-fed obese animal models (Fig.2). Further evidences showed that this was due to PBG’s ability to improve intestinal barrier integrity and thus decrease leakage of microbial LPS from gut into circulation.

The modulation of the gut microbiotic composition offers a new avenue for the treatment of obesity and metabolic disorders (18). PBG is identified for the first time as an anti-obesity agent. The transfer of feces from PBG-treated mice protected the recipient HFD-fed mice from obesity, suggesting a vital role of gut microbiota in PBG’s anti-obesity effect. Unlike other established prebiotics such as dietary polyphenols (29), inulin (30), and oat-derived β-glucan (31), PBG didn’t restore HFD-induced increase in the *Firmicutes* to *Bacteroidetes* ratio. PBG markedly increased the abundance of *A. muciniphila*, a mucin-degrading bacterium that could reverse HFD-induced obesity (32). PBG didn’t boost the growth of *A. muciniphila*, either in pure cultures or gut-simulators. By use of a cocktail of antibiotics, microbiota-depleted mice were generated. Data from microbiota-depleted mice indicated that PBG created an intestinal microenvironment favorable for *A. muciniphila* colonization. The genome of *A. muciniphila* contains a large proportion of genes encoding secreted proteins (567 of the 2,176 open reading frames), 61 of which have been assigned protease, sugar hydrolase, sialidase, or sulfatase activities, suggesting specialization in mucus utilization and adaptation to the gut environment (33). Given recent evidence indicated that *A. muciniphila* colonization was mediated by host immune status (34, 35), it will be intriguing to clarify the relationship between richness of *A. muciniphila* and PBG-induced immunomodulation in the future.

Transcriptome analysis of PBG-induced differentially expressed genes in colon tissues indicated that PBG had a direct pronounced effect on host lipid metabolism. These finding implied that PBG might reduce fat accumulation through a microbiota-dependent or -independent manner.

In summary, we demonstrated that PBG, one representative of *C. versicolor* polysaccharopeptide originally used as an adjunct therapy for cancers could be an anti-obesity agent through modulating gut microbiota and regulating host metabolism. Overweight and obesity are associated with at least 13 different types of cancer. These cancers make up 40% of all cancers diagnosed (36, 37). A variety of biological mechanisms involving the adipocyte has been implicated in tumorigenesis. The previous consensus on the anti-cancer effect and the current knowledge in the anti-obesity effect of *C. versicolor* polysaccharopeptides might make them favorable to those patients suffered from obesity-associated cancers.

## Methods

### Murine

C57BL/6J female mice were purchased from the Fourth Military Medical University (Xi’an, Shaanxi, China). After one week for accommodation, mice were randomly distributed into different groups, fed with either a standard chow diet (10 kcal % Fat, D12492) or a high-fat diet (60 kcal % Fat, D12450J). Animal procedures were approved by the Animal Ethics Committee in Northwest A & F University, China.

### Preparation of protein-bound β-glucan

The *Coriolus versicolor* powder was subjected to hot water extraction and alcohol precipitation. The resulting precipitates were dissolved in water and dialyzed (cutting MW at 8000 Da). The non-dialysates were concentrated and the supernatant was applied to a DEAE-Fast-Flow column (10 i.d. × 60 cm) with stepwise distilled H_2_O and 0.5 M NaCl solution. The 0.5 M NaCl eluent, designated as PBG was dialyzed and lyophilized. The structural information was shown in Table S2.

### Body composition analysis

Body composition was measured with the nuclear magnetic resonance system using a Body Composition Analyzer MiniQMR23-060H-I (Niumag, Shanghai, China). Body fat and lean mass were determined in live conscious mice with adlibitum access to chow as previously described (38).

### H&E Staining

Liver was fixed in 10% buffered formalin at room temperature before embedding in paraffin. Histological assessment of H&E sections was performed in a blinded fashion by a pathologist using a previously described scoring system (39).

### Cytokine measurements

IL-1β, IL-6, IL-10 and TNF-α protein levels in serum were measured using commercial Bio-Plex Pro mouse kits (BIO-RAD, USA).

### Biochemical analysis

Serum leptin, adiponectin, and LPS were measured by quantification assay kits (Cusabio, China). Triglyceride and total BA were assayed using commercial detection kits (Nanjing Jiancheng Chemical Industrial Co. Ltd, China).

### Real-Time PCR

Samples of total RNA from colon tissue was isolated using the TRIZOL solution (TransGen, Beijing, China). Quantitative real-time reverse-transcription PCR (qRT-PCR) was performed in triplicate on a QuantStudio™ 6 Flex Real-Time PCR (Life Technologies, Singapore). The primers used are shown in Table S3 (40, 41).

### Gut microbiota analysis

Stool samples were snap-frozen in liquid nitrogen before storage at −80°C. Total bacterial DNA was extracted using a fecal DNA isolation kit (MoBio Laboratories, USA) according to the manufacturer’s protocol. The 16S rRNA gene comprising V3–V4 regions was amplified using common primer pair and the microbial diversity analysis was performed as described (42).

Briefly, the raw sequences were first quality-controlled using QIIMEwith default parameters, then demultiplexed and clustered into species-level (97% similarity) operational taxonomic units (OTUs). OUT generation is based on GreenGene’s database and the reference-based method with SortMeRNA. Strain composition analysis, alpha diversity analysis and beta diversity analysis were also performed using QIIME. Discriminative taxa were determined using LEfSe (LDA Effect Size, http://huttenhower.sph.harvard.edu/galaxy/).

### *In vitro* gut simulator

A three compartment dynamic *in vitro* human intestinal tract model (SHIME) was used to study the effects of PBG on a stabilized gut microbial community in a controlled *in vitro* setting (43).

### *A. muciniphila* colonization

C57Bl/6J female mice (4 weeks old) were fed with HFD and treated with an antibiotic cocktail for 4 weeks as described (44). The microbiota-depleted mice were daily supplementated with or without PBG (400 mg/kg) for another month in the presence of antibiotics. Mice were lavaged with *A. muciniphila* (1.5 × 10^8^ cfu) continuously for 3 days as previously described (32). Fecal samples were collected 24 h after each lavage and *Akkermansia muciniphila* in the feces was quantified by qPCR using the universal 16S rRNA gene primers (shown in Table S3).

### Fecal transplantation

Fecal transplantation was performed according to a previous study (45). The donor mice (4-week-old, n=10 per group) were fed with Chow, (PBG+Chow), (PBG+HFD), or HFD for 4 weeks. Stools from donor mice of each diet group were subsequently collected and pooled. The transplant material was prepared as resuspension of 100 mg of stools from each diet group in 1 ml of sterile saline, and centrifugation at 800g for 3 min to obtain the supernatant. The recipient mice (8-week-old, n=6-8 per transplant group) were fed with HFD and orally treated with 200μl of fresh transplant material daily, which was prepared on the same day within 10 min of transplantation. After 8 weeks of treatment, the recipient mice were killed for subsequent analysis.

### Transcriptomic analysis

HFD-fed mice were treated daily with a cocktail of antibiotics for one month, followed by supplementation with or without PBG (400 mg/kg) for another month in the presence of antibiotics (46). RNA extracted from colon tissues was performed via a paired-end 125 cycle rapid run on the Illumina HiSeq. 2500. The clean reads that were filtered from the raw reads were mapped to mouse (C57BL/6 strain) reference genome (GRCm38) using Tophat2 (47) software. FPKM values were used to estimate gene expression by use of the Cufflinks software (26). The DESeq (48), Kyoto Encyclopaedia of Genes and Genomes (KEGG), and gene ontology (GO) terms were determined via protein database by BLASTX (49).

### Statistical analysis

All data of experiment were shown as means± standard error of mean (S.E.M.) Data sets that involved more than two groups were assessed by one-way ANOVA followed by Dunnett’s multiple comparisons test and unpaired two-tailed Student’s t-test. 16S rRNA gene sequence analysis was assessed using Paired Wilcoxon rank-sum test. A *p* value of 0.05 was considered statistically significant based on ANOVA statistical analysis by Graphpad 7.0 and the R programming language. Data of RNA-Sequence are presented as mean FPKM ± S.E.M.. Differences between PBG and Blank mice that were evaluated FDR (*p* < 0.05) by using Benjamini-Hochberg method.

## Acknowledgement

This work was supported by the National Natural Science Foundation of China (NSFC) (31570799), and Fundamental Research Funds for the Central Universities, Northwest A&F University (2452017026).

